# Quantification of anti-parasite and anti-disease immunity to malaria as a function of age and exposure

**DOI:** 10.1101/191197

**Authors:** Isabel Rodriguez-Barraquer, Emmanuel Arinaitwe, Prasanna Jagannathan, Moses R Kamya, Philip J. Rosenthal, John Rek, Grant Dorsey, Joaniter Nankabirwa, Sarah G. Staedke, Maxwell Kilama, Chris Drakeley, Isaac Ssewanyana, David L Smith, Bryan Greenhouse

## Abstract

Malaria immunity is complex and multi-faceted, and fundamental gaps remain in our understanding of how it develops. Here, we use detailed clinical and entomological data from three parallel cohort studies conducted across the malaria transmission spectrum in Uganda to quantify the development of immunity against symptomatic *Plasmodium falciparum* as a function of age and transmission intensity. We focus on: anti-parasite immunity (i.e; ability to control parasite densities) and anti-disease immunity (i.e; ability to tolerate higher parasite densities without fever). Our findings suggest a strong effect of age on both types of immunity, that remains significant after adjusting for cumulative exposure. They also show a non-linear effect of transmission intensity, where children experiencing the lowest transmission appear to develop immunity faster than those experiencing higher transmission. These findings illustrate how anti-parasite and anti-disease immunity develop in parallel, reducing the probability of experiencing symptomatic malaria upon each subsequent *P. falciparum* infection.

## Introduction

The last decades have seen substantial declines in malaria transmission in sub-Saharan Africa that are largely attributable to increased access to effective control measures, including insecticide-treated bednets, indoor residual spraying of insecticide and artemisinin-based combination therapy(1,2). In settings where transmission has been low, increased access to effective control interventions opens the possibility for malaria elimination. In highly endemic settings, however, there are concerns around the potential impact of failing to sustain interventions that reduce but do not stop transmission. Short-term decreases in malaria incidence due to reductions in transmission could be offset over time by reductions in population immunity to malaria resulting from lower exposure to parasites (3–5).

Gradual acquisition of immunity against symptomatic malaria is a key driver of the epidemiology of malaria in endemic settings, where the incidence of disease typically peaks in early childhood and then declines, while the prevalence of detectable asymptomatic parasitemia increases throughout childhood before declining in adulthood (6–12). While these epidemiologic patterns have been described across the transmission spectrum, there are still many fundamental gaps in our understanding of the factors driving the development of immunity, and of the independent roles of age and repeated infection. One reason it has been challenging to study immunity to malaria is that there are currently no reliable and quantifiable immune correlates of protection that can be measured in epidemiological studies. In addition, there are few available datasets that include both detailed clinical data and independent metrics of exposure at the individual level.

Here, we use data from three parallel cohort studies conducted across the spectrum of malaria transmission in Uganda to model and quantify the development of immunity against symptomatic malaria as a function of transmission intensity and age. A key strength of these studies is that they involved detailed clinical and entomological surveillance of all study households. We focus on two specific types of immunity: anti-parasite immunity (i.e; the ability to control parasite densities upon infection) and anti-disease immunity (i.e; the ability to tolerate higher parasite infections without developing objective fever), as they have been described as independent components of clinical immunity (13).

## Results

The three cohorts enrolled a total of 1021 children aged 6 months to 10 years from 331 randomly chosen households across the three study sites. This analysis was limited to data from 773 children who experienced at least one patent *P. falciparum* infection between August 2011 and November 2014. Table 1 summarizes the general characteristics of the participants included in this analysis.

**Table 1:** Characteristics of the study participants

| Characteristic | Nagongera | Kihihi | Walukuba |
| --- | --- | --- | --- |
| Number of households | 106 | 100 | 76 |
| Number of children | 329 | 305 | 139 |
| Female, *n* (%) | 151(46) | 150 (49) | 66 (47) |
| Mean age at enrollment, years (*sd*) | 4.4 (2.7) | 4.6(2.6) | 4.3 (2.6) |
| Mean follow up time, months (range) | 23.5 (0, 38.8) | 24.4 (0.8, 38.8) | 22.1 (2.3, 3.9) |
| *Symptomatic malaria* |  |  |  |
| Symptomatic Malaria episodes, *n* | 2447 | 1555 | 207 |
| Median number of symptomatic malaria episodes/child, *n* (range) | 6 (0, 29) | 4 (0, 30) | 1 (0, 12) |
| Median incidence of symptomatic malaria episodes ppy (range) | 2.6(0, 10) | 1.6 (0, 15.2) | 0.6 (0, 5.1) |
| *Asymptomatic parasitemia* |  |  |  |
| Asymptomatic parasitemia episodes, *n* | 955 | 331 | 145 |
| Median number of asymptomatic parasitemia episode/child, *n* (range) | 2 (0, 12) | 0 (0, 11) | 1 (0, 10) |
| Median prevalence of asymptomatic parasitemia (range) | 0.12 (0.07-0.17) | 0.05 (0.02-0.10) | 0.07 (0.03-0.11) |
| *Household malaria exposure* |  |  |  |
| Household aEIR, median (range) | 51 (10-582) | 7.7 (3.6-47) | 2.1 (1.5-8.1) |

Participants living in Nagongera experienced the highest incidences of symptomatic malaria (median 2.6 episodes per person year), followed by those living in Kihihi (median 1.6 episodes per person year) and Walukuba (median 0.6 episodes per person year) (Table 1 and Figure 1). These incidences were consistent with results from monthly entomological surveys conducted in all cohort households, with significantly higher annual entomological inoculation rates (aEIR) recorded in Nagongera (median 51 infectious bites per year, range 10-582) as compared to Kihihi (median 8 infectious bites per year, range 4-47) and Walukuba (median 2 infectious bites per year, range 1-8). Interestingly, prevalence of asymptomatic parasitemia did not follow this same relationship; the prevalence of asymptomatic parasitemia was highest in Nagongera, and prevalences in the lower transmission sites were similar.

**Figure 1:**
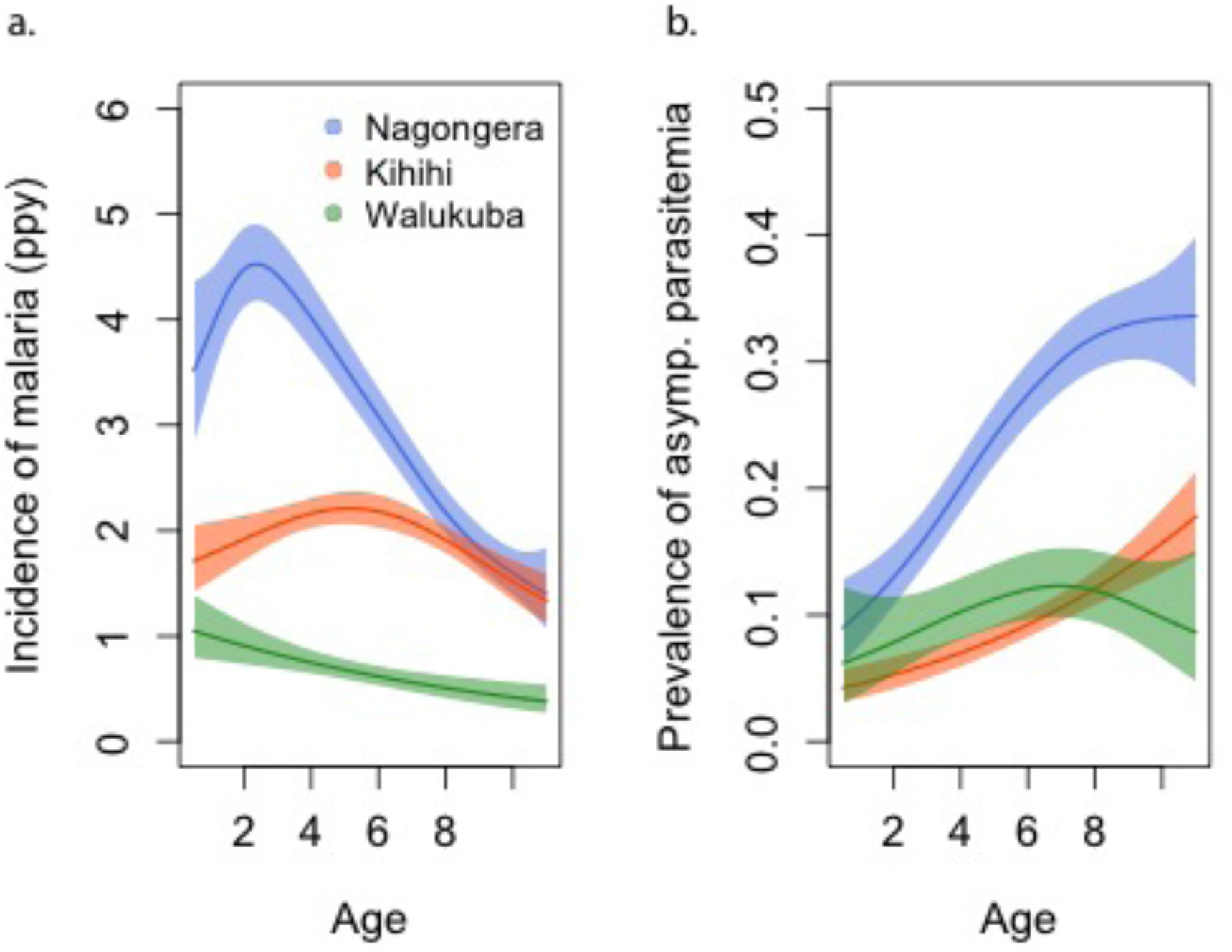
Incidence of malaria **(a)** and prevalence of asymptomatic parasitemia **(b)** in the three study sites as a function of age. Shaded areas represent 95% confidence bounds.

### aEIR as a metric of individual exposure

To assess whether entomological metrics were a good indicator of individual exposure to *P. falciparum*, we correlated the measured annual EIRs (aEIR) for each household (Figure 2a) with estimates of the average individual hazard of infection (Figure 2b). Individual hazards were estimated by fitting time-to-event models to the incidence data from each site. We found a significant correlation between these two independent metrics of exposure across sites (R^2^ = 0.38, p<0.001). While aEIR explained less of the variance between individuals within each site, the correlation was still significant for both Nagongera (R^2^ = 0.03, p=0.005) and Kanungu (R^2^ = 0.12, p<0.001).

**Figure 2:**
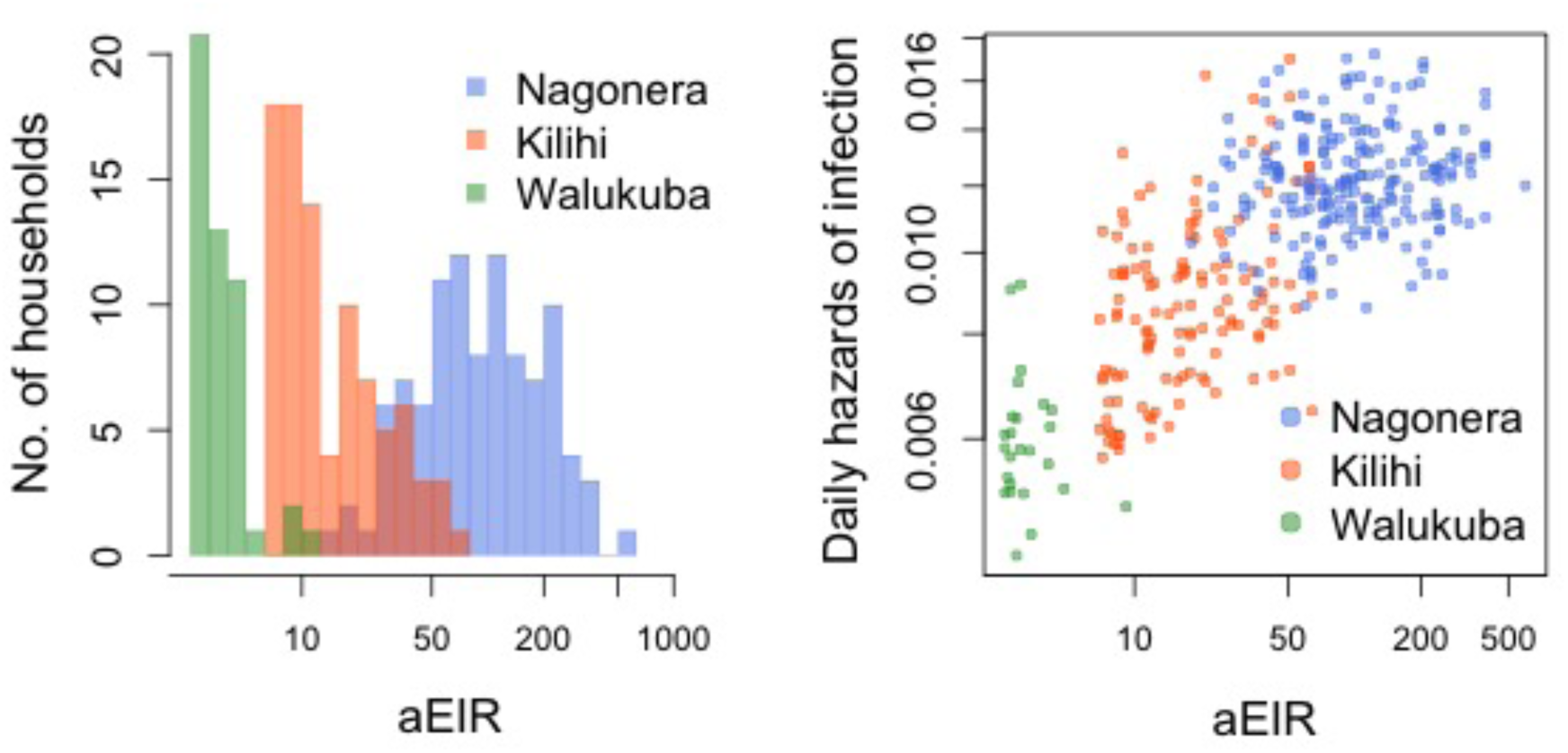
**a.** Distribution of the average annual entomological inoculation rate (aEIR) experienced by the study households in the three study sites. **b.** Correlation between the measured aEIRs and the estimated individual hazards of infection.

### Anti-parasite immunity

Parasite densities developed upon infection decreased with increasing age in all settings and for both symptomatic (passive detection) and asymptomatic (detected during routine visits) infections. Despite the large variability in parasite densities recorded within and between individuals, this trend is evident in the raw data (Figure 3a). A trend towards lower parasite densities was also observed among individuals living in settings with higher aEIRs (Nagongera), as compared to settings with lower aEIR (Kihihi and Walukuba).

**Figure 3:**
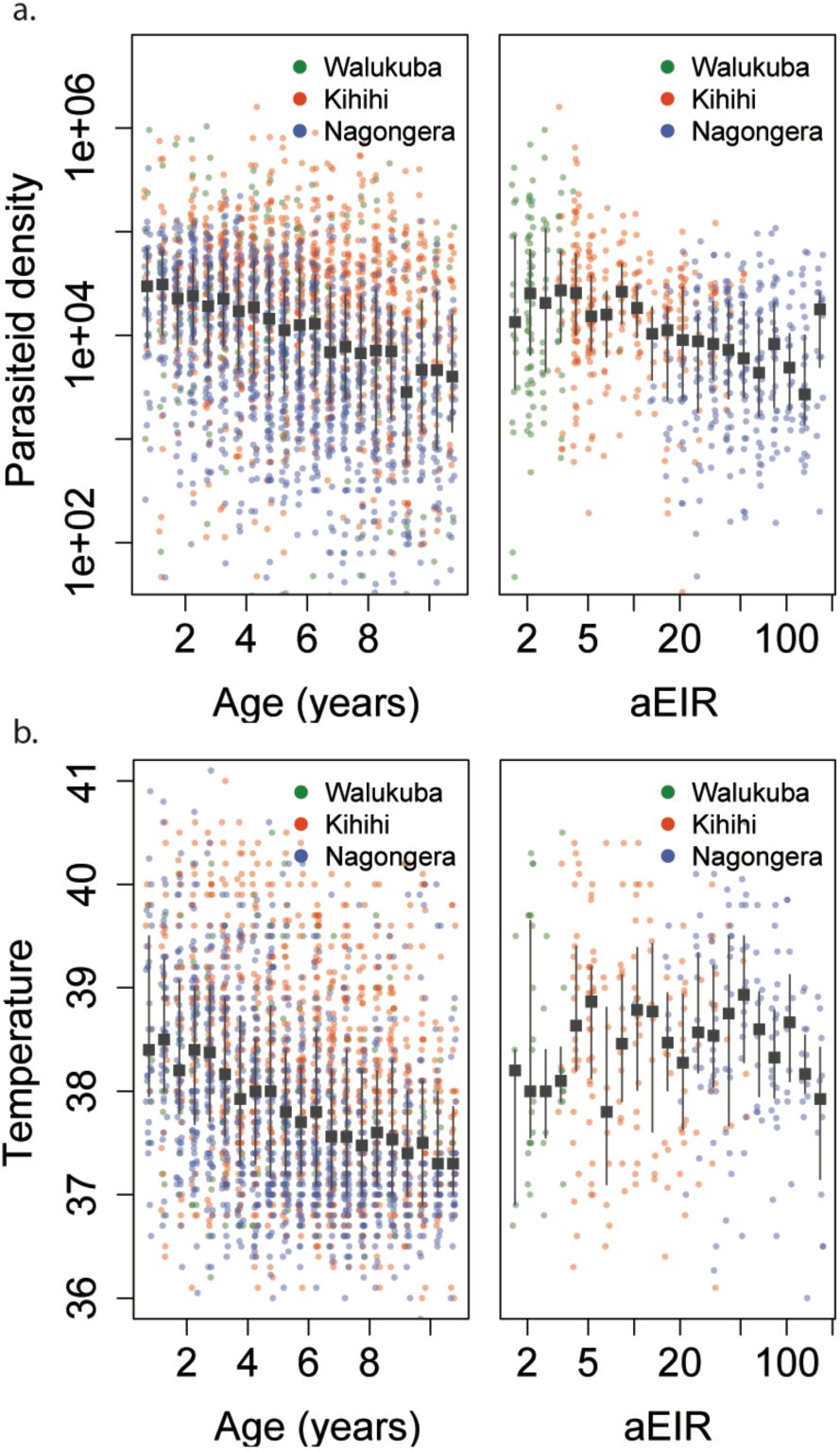
Trends in parasite densities **(a)** recorded during symptomatic (passive surveillance) infections and routine (active surveillance) visits as a function of age (left) and aEIR (right); and trends in the objective temperature **(b)** recorded during visits in which participants were found to have a parasite density between 50,000 parasites/μL and 200,000 parasites/μL, as a function of age (left) and aEIR (right). Each point represents a measurement obtained during a study visit. The median and interquartile range are shown in black.

We considered multiple candidate models to describe the association between parasite density, age and aEIR (supplementary material). Models allowing smooth (non-linear) relationships with aEIR best fit the data. Models allowing for two-way interactions between age and aEIR also outperformed models that didn’t include interactions.

In moderate and high transmission settings (households with aEIR >5), increasing age and increasing exposure were independently associated with decreases in the parasite densities. On average, parasite densities decreased by a factor of 0.76 (95%CI 0.74-0.77) for each additional year of age and by a factor of 0.73 (95%CI 0.70-0.76) for each two-fold increase in the aEIR. The relationship was less evident for the lower transmission households (aEIR<5). In these settings, there continued to be a decreasing association with age, but the expected parasite densities at any given age were equal or lower to those observed in the higher exposure (aEIR>10) settings.

Figures 4a and 5a present the predicted parasite densities, as a function of age and aEIR, according to the best fitting model. While an individual aged 1 year exposed to an aEIR of 10 is expected to develop a parasite density of 14610 parasites/μL (95%CI 5924– 36031 parasites/μL) upon infection, the expected parasite density goes down to 3237 parasites/μL (95%CI 1381–7586 parasites/μL) by age 10 years. In contrast, the expected parasite density in an individual living in a setting with aEIR of 150 will be similar 13071 parasites/μL (95%CI 5256– 32503 parasites/μL) at age 1 year, but significantly lower by age 10 years (999 parasites/μL (95%CI 398–2508 parasites/μL)).

**Figure 4:**
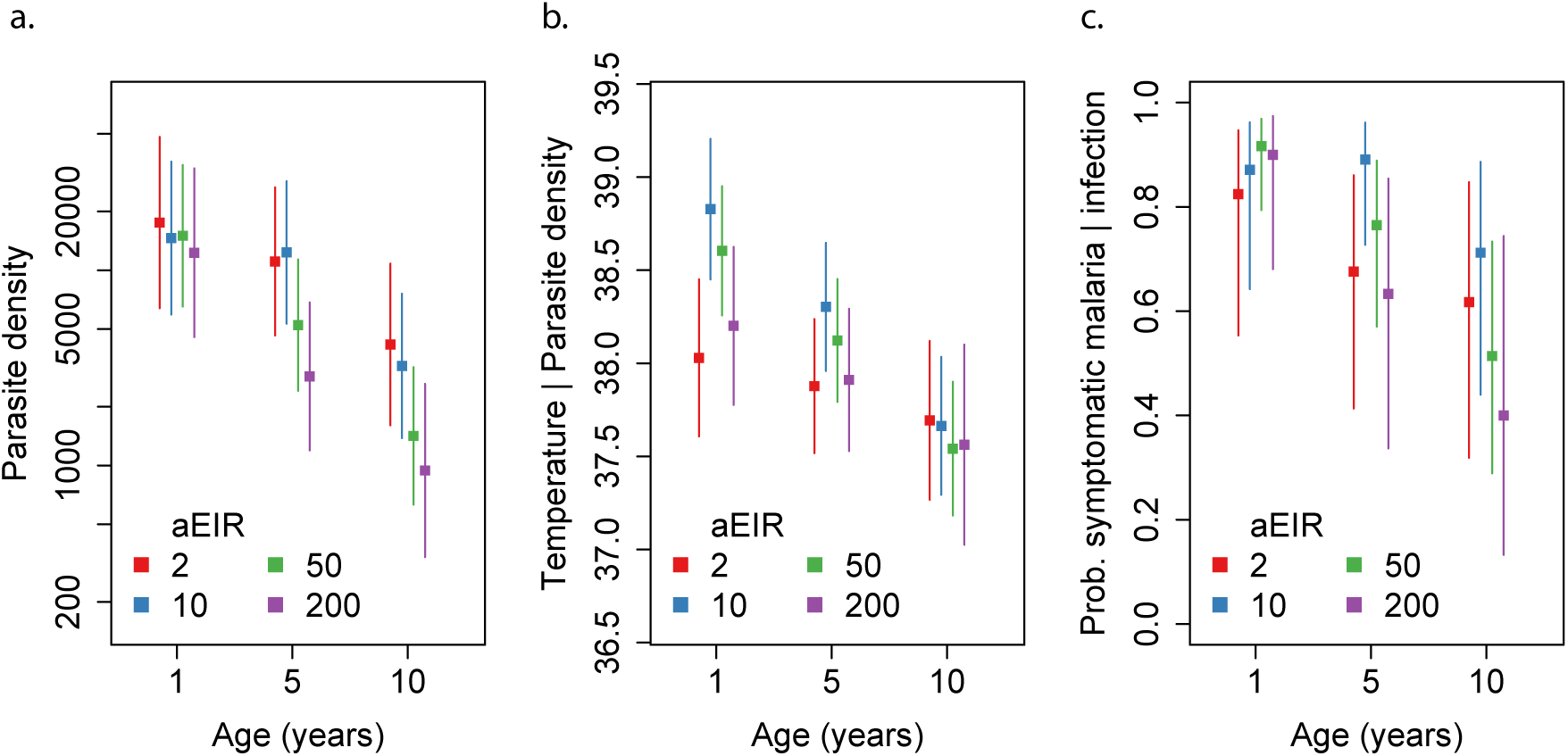
Results from models quantifying anti-parasite immunity **(a),** anti-disease immunity **(b)** and overall immunity against symptomatic malaria **(c).** Each plot shows, for specific ages and aEIRs, the expected parasite density (/μL) **(a)**, objective temperature given a density of 40,000 parasites/μL **(b)** and the probability of developing symptomatic malaria upon infection **(c),** estimated using the best fitting model. 95% confidence intervals of the estimates are also shown.

To test whether the observed associations with age could be explained by the cumulative exposure over a life time, we also fit models where, instead of adjusting for the aEIR, we adjusted for the cumulative number of infectious bites (i.e. the product of age and aEIR). Results from these models are consistent with a smaller, yet independent effect of age on the development of anti-parasite immunity; for any given level of cumulative exposure, each additional year of life was associated with decreases in parasite densities by a factor of 0.82 (95%CI 0.81-0.85).

### Anti-disease immunity

We define anti-disease immunity as the ability to tolerate a given parasite density without developing objective fever. Thus, we were interested in modeling temperatures recorded at specific parasite densities, as a function of age and aEIR. Similar to the models characterizing anti-parasite immunity, models including smooth effects and interactions outperformed simpler models.

As expected, we found a strong association between parasite densities and objective temperature (Figure S1). Increases in parasite densities above 1000 parasites/μL were associated with higher expected temperatures across ages and transmission settings. In addition, we found a negative association between objective temperature at a given parasite density and age (Figures 3b, 4b and 6). In moderate and high transmission settings (aEIR>5), the objective temperature at a given parasite density decreased on average by 0.08 °C (95%CI 0.07-0.10 °C) for each additional year of life. Thus, while the39.2 expected temperature for a child aged 1 year living in a setting with aEIR of 10 with a parasite density of 40,000 would be 38.8 °C (95% CI 38.5-°C), the expected38.0 temperature would decrease to 37.6 °C (95% CI 37.3-°C) if the same child experienced the infection at age 10 years (Figure 6). This association was similar and remained significant even when adjusting for cumulative exposure and for the differences in incidence of non-malarial fever across age-groups (Figure S3).

Similar to the anti-parasite immunity results described above, the observed association between exposure level and anti-disease immunity was less evident than the association with age. For moderate and high transmission settings (aEIR 5 to 300) there was a clear negative association between objective temperature at a given parasite density and aEIR. The objective temperature decreased by 0.07 °C (95%CI 0.05-0.10 °C) for each two-fold increase in aEIR. However, the relationship did not follow this trend for lower transmission settings. Children living in the lowest transmission settings (aEIR 1 to 5) appeared to tolerate higher parasite densities than children living in moderate transmission settings (aEIR 5 to 10).

As an alternative way to characterize anti-disease immunity, we used our best fitting model to predict the fever threshold, defined as the minimum parasite density associated with objective fever (temperature> 38 °C), across levels of age and aEIR (Figure 5b). Results from this analysis show that, for settings with moderate and high transmission (aEIR>5), the fever threshold increases both with age and increasing exposure. Thus, while a 1 year old child living in a setting with aEIR of 10 presenting with a parasite density as low as 3747 parasites/μL (95%CI 777-11129 parasites/μL) will be expected to be febrile, children older than 6 years of age exposed to very high transmission (aEIR 150) might be afebrile even with parasite densities higher than 60000 parasites/μL.

**Figure 5.**
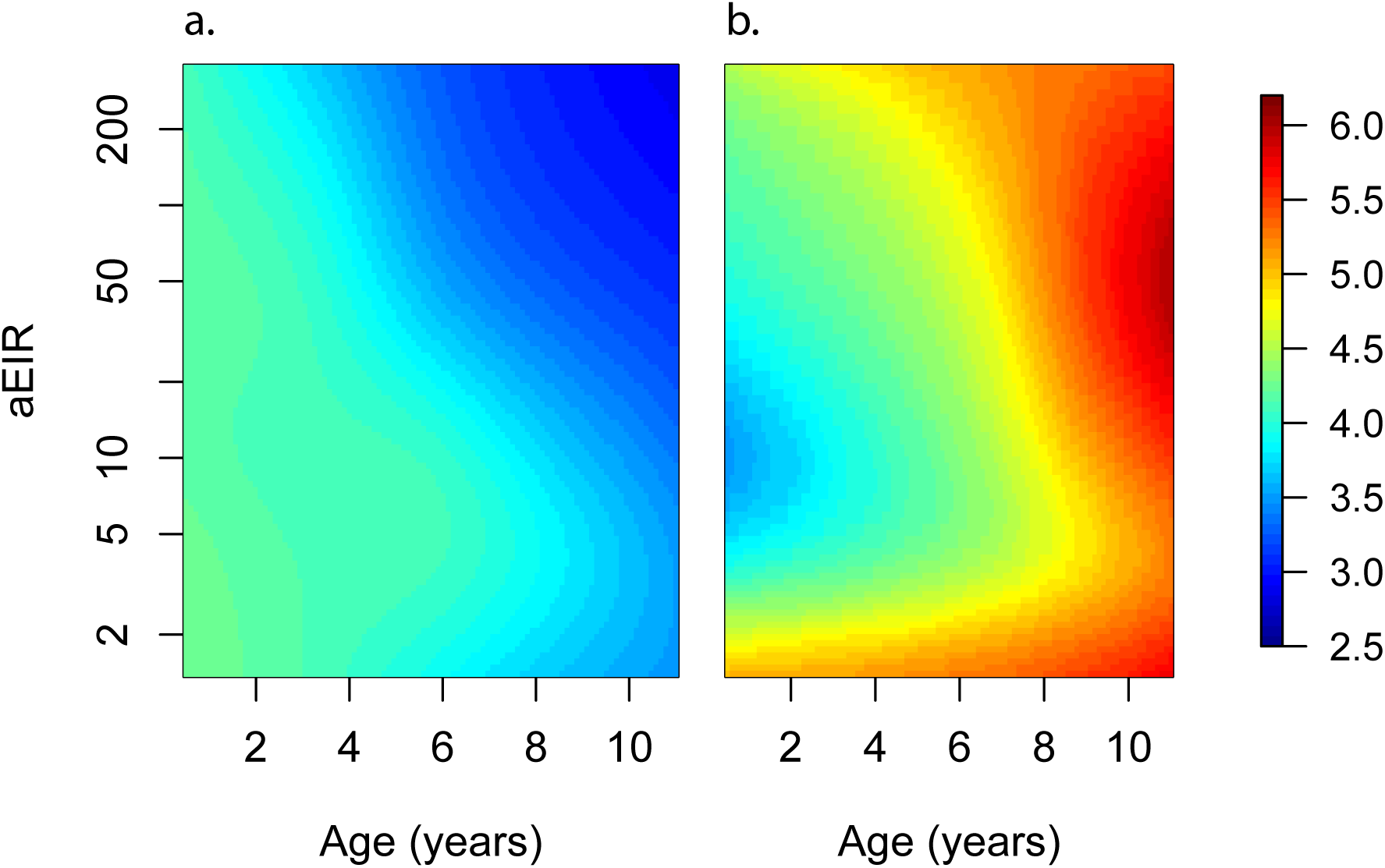
Results of models quantifying anti-parasite (**a**) and anti-disease immunity (**b**). These results are similar to those presented in Figure 4, but for the full range of ages and aEIRs included in the data. Panel **a.** shows expected parasite densities (log 10) after infection at different ages and levels of exposure (aEIR). Panel **b.** shows the expected fever thresholds (parasite densities required to develop a temperature 38°C or greater). Variance estimates for these plots are presented in the supplementary materials.

**Figure 6.**
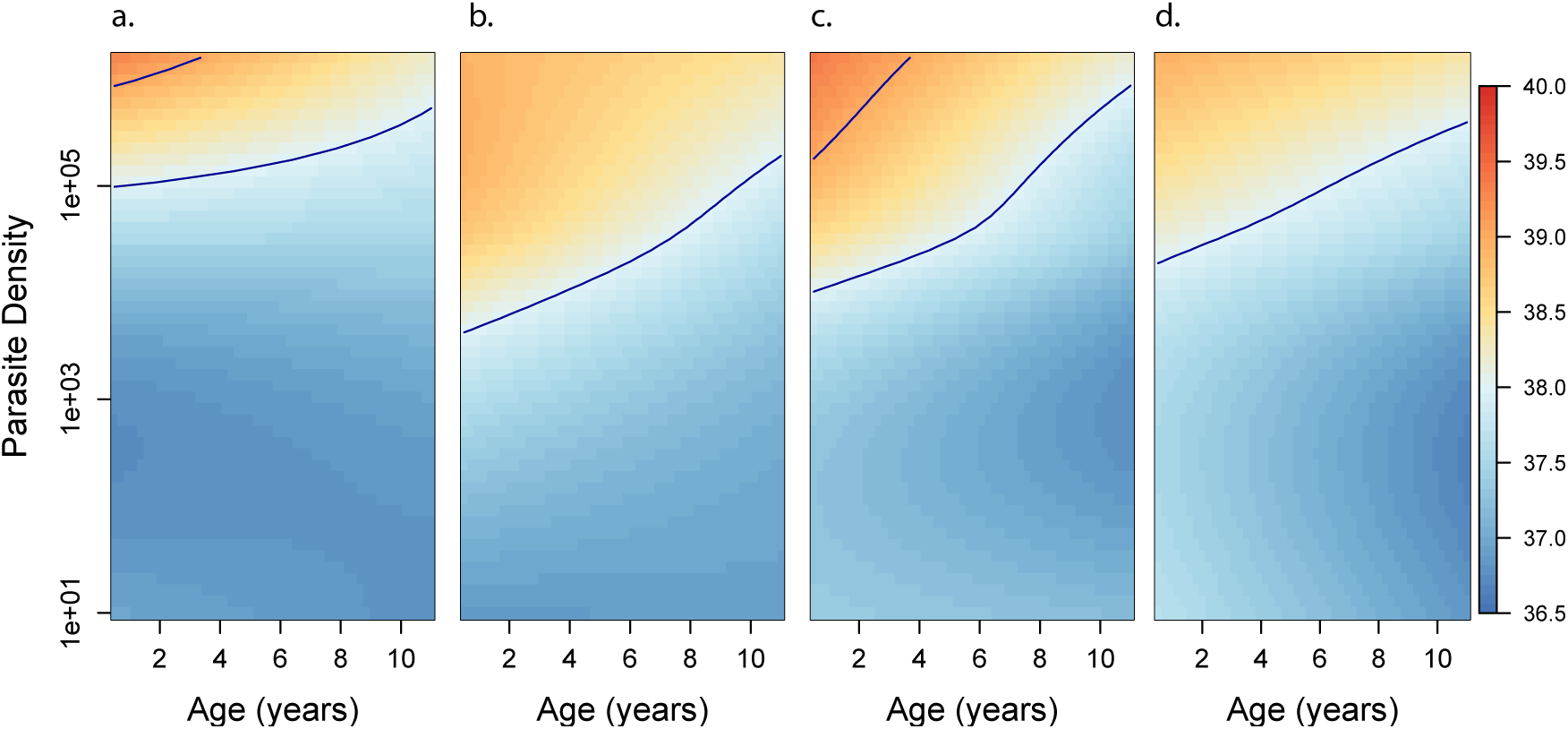
Results of the best model quantifying anti-disease immunity. Each panel shows how the expected objective temperature (°C) varies as a function of age and parasite density, for different transmission settings. **a.** aEIR=2; **b.** aEIR=10; **c.** aEIR=50; **d.** aEIR=200. Contours indicating the fever threshold (38°C) and 39°C are also shown. Variance estimates for these plots are presented in the supplementary materials.

### Overall immunity against symptomatic malaria

Finally, to characterize the association between age and aEIR on the overall risk of developing symptomatic malaria upon infection (i.e.; the combined effect of anti-parasite and anti-disease immunity), we fit a series of models where the outcome of each independent observed infection (i.e.; symptomatic malaria or asymptomatic parasitemia) was modeled as a function of age and aEIR.

Results from this analysis are consistent with results from the anti-parasite and anti-disease models (Figure 7). While young children living in low transmission settings (aEIR=5) are expected to develop symptomatic malaria in most their infections, the probability that an infection results in symptomatic malaria decreases as a function of age and exposure. The expected probability of symptomatic disease for a child aged 1 year living in a setting with aEIR of 50 is 0.92 (95%CI 0.79-0.97), but it decreases to 0.51 (95%CI 0.29-0.73) by age 10 years.

**Figure 7.**
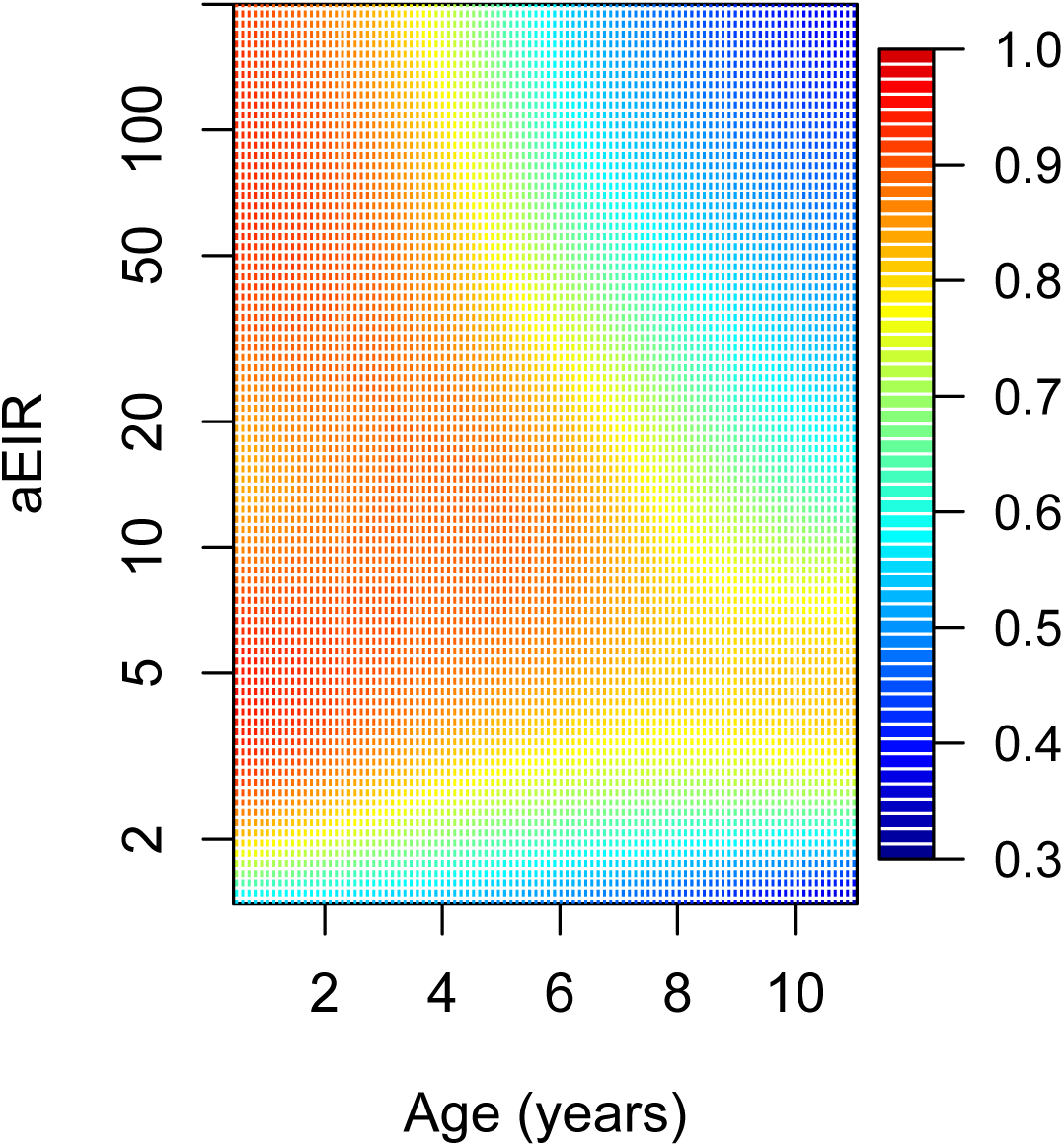
Results of the best model exploring overall immunity against symptomatic malaria, as a function of age and exposure (aEIR). Colors represent the probability of developing symptomatic malaria upon infection. Variance estimates for these plots are presented in the supplementary materials.

### Sensitivity analyses

Our main analyses include data from all visits regardless of their type (routine vs passive case detection). Thus, the expected values modeled here may be biased by the frequency of active vs passive episodes detected. In particular, it is possible that we have under-sampled the instances of asymptomatic disease, and thus, our estimates of the expected parasite densities may be an over-estimate of those present in the population. To address this, we performed sensitivity analyses where we up-weighted the episodes of asymptomatic parasitemia, to account for potentially unobserved asymptomatic infections. Results from these analyses were qualitatively identical to the main analysis reported here and are presented in the supplementary material (Figure S4).

To explore whether differences in the prevalence of certain host genetic polymorphisms between sites could be driving some of our findings, we also performed sensitivity analyses limiting the dataset to those subjects without the sickle hemoglobin mutation (β globin E6V), known to protect against malaria (17,18). Even though the sample size of these analyses was significantly smaller, results were unchanged qualitatively (Figure S5). Similarly, restricting the dataset to children without two other known polymorphisms (the α-thalassemia 3.7 kb deletion or glucose-6-phosphate dehydrogenase deficiency caused by the common African variant (G6PD A-)), had little effect on the results.

## Discussion

Our findings illustrate how anti-parasite and anti-disease immunity develop gradually and in parallel, complementing each other in reducing the probability of experiencing symptomatic disease upon infection with *P. falciparum*. While anti-parasite immunity acts to restrict the parasite densities that develop upon each subsequent infection, anti-disease immunity increases the tolerance to high parasite densities. Thus, older children are less likely to develop symptomatic malaria upon infection both because they tolerate parasite densities better without developing fever, and because they are less likely to develop high parasite densities.

Our results indicate independent effects of age on the acquisition of both anti-parasite and anti-disease immunity. These independent age effects may reflect maturation of the immune system as well as other physiological changes that decrease the propensity to fever (13,19). Furthermore, our findings are consistent with independent effects of transmission intensity on the acquisition of these two types of immunity. While the results obtained for moderate and high transmission settings (aEIR >5) are consistent and expected, and suggest that immunity develops faster in settings where individuals get infected by *P. falciparum* more often, the results obtained for the lowest transmission settings are harder to reconcile. These results were largely driven by observations collected in the Walukuba site, and as such it is possible that site-specific characteristics may have driven them. Walukuba was previously a relatively high transmission rural area, but substantial decreases in transmission intensity have been observed since 2011, likely due to urbanization. While our sensitivity analyses suggested that differences in the prevalence of three well characterized host-genetic polymorphisms between sites do not explain these discrepant results, it is still possible that other unmeasured site-specific characteristics may have driven them. A lower parasite diversity in Walukuba, for example, could cause this difference, as developing an effective immune response against fewer parasite strains may be much easier than developing immunity against a more diverse pool (20,21). Testing this hypothesis would require careful characterization of the complexity and diversity of infections in each of our cohort settings.

While site specific characteristics may underlie the observed high levels of clinical immunity against malaria in the low transmission setting, it is also possible that this finding reflects biologically relevant differences in how immunity against malaria develops. For example, it has been hypothesized that immunity may develop optimally in individuals that are exposed at a low rate, and that more frequent infections may interfere with the development of robust immune responses (22,23). Answering this question will require further detailed studies across transmission settings, with careful characterization of both exposure and infection outcomes.

There are several limitations to this study. With a study design including routine visits every 3 months, we are likely to have missed several asymptomatic infections, particularly in the moderate and high transmission settings. Moreover, since infections were detected using microscopy, we were unable to detect sub-patent infections, and we lack knowledge about the genetic complexity of each infection. While it is possible that the expected values modeled here (expected parasite density and fever threshold) were biased by these sources of measurement error, sensitivity analyses suggest that the relationships observed were robust. Secondly, while we found an independent association between the average household aEIR and both anti-parasite and anti-disease immunity, it is not clear that this is the most relevant metric of exposure for the development of clinical immunity to malaria. Alternative metrics such as the number of discrete infections, the number of “strains” seen or the total parasite-positive time might be more relevant, but require the collection of additional data, including more frequent sampling. Finally, while this study provides very detailed insight into how two types of clinical immunity to malaria develop in endemic settings as a function of age and repeated exposure, it says nothing about the duration of immunity.

Prior studies have tried to model the processes driving acquisition of clinical immunity against malaria. However, these models have been generally informed by aggregated epidemiological data (age-incidence and age-prevalence) which limits their capacity to isolate the contributions of age and repeated exposure (24–26). Our results quantify how anti-parasite and anti-disease immunity develop in children across the malaria transmission spectrum, and they support strong a strong independent effect of age and a perhaps paradoxical effect of exposure. The methods proposed here to model anti-parasite and anti-disease immunity may also provide a framework to select individuals with immune and non-immune phenotypes for evaluations of immune correlates of protection.

## Methods

### Ethics Statement

The study protocol was reviewed and approved by the Makerere University School of Medicine Research and Ethics Committee, the Uganda National Council for Science and Technology, the London School of Hygiene & Tropical Medicine Ethics Committee, the Durham University School of Biological and Biomedical Sciences Ethics Committee, and the University of California, San Francisco, Committee on Human Research. All parents/guardians were asked to provide written informed consent at the time of enrollment.

### Data

We used data from three parallel cohort studies conducted in Uganda in sub-counties chosen to represent varied malaria transmission(14). Walukuba, in Jinja district, is a peri-urban area near Lake Victoria that has the lowest transmission among the three (annual entomological inoculation rate (aEIR) estimated to be 2.8(14)). Kihihi, in Kanungu district, is a rural area in southwestern Uganda characterized by moderate transmission (aEIR=32). Nagongera, Tororo district, is a rural area in southeastern Uganda with the highest transmission (aEIR=310)(14,15). Details on how the study households and participants were selected has been described elsewhere(14). Briefly, all households were enumerated, and then approximately 100 households were selected at random from each site. Between August and September 2011, all children from these households aged between 6 months and 10 years who met eligibility criteria were invited to participate. As the cohorts were dynamic, additional children from participating households were invited to participate if they became eligible while the study was ongoing. Unless participants were withdrawn from the study either voluntarily or because they failed to comply with study visits, they were followed-up until they reached 11 years of age. Children from 31 randomly selected additional households were enrolled between August and October 2013 to replace households in which all study participants had been withdrawn. For this analysis, we used data collected from visits between August 2011 and November 2014. The studies included passive and active follow-up of participants. Parents/guardians were encouraged to bring their children to designated study clinics for any illness. All medical care was provided free of charge, and participants were reimbursed for transportation costs. All children who reported fever in the previous 24 hours or were febrile at the time of the visit (tympanic temperature > 38.0°C) were tested for malaria infection with a thick blood smear. Light microscopy was performed by an experienced laboratory technician who was not involved in direct patient care and verified by a second technician. Parasite density was calculated by counting the number of asexual parasites per 200 leukocytes (or per 500 leukocytes, if the count was <10 asexual parasites/200 leukocytes), assuming a leukocyte count of 8,000/μl. A blood smear was considered negative when no asexual parasites were found after examination of 100 high power fields.

If the smear was positive, the patient was diagnosed with symptomatic malaria and received treatment with artemether-lumefantrine (AL), the recommended first-line treatment in Uganda. Episodes of complicated or recurrent malaria occurring within 14 days of therapy were treated with quinine. In addition, routine evaluations were performed every three months, including testing for asymptomatic parasitemia using thick blood smears.

Entomological surveys were also conducted every month at all study households. During these surveys, mosquitoes were collected using miniature CDC light traps (Model 512; John W. Hock Company). Established taxonomic keys were used to identify female *Anopheles* mosquitoes. Individual mosquitoes were tested for sporozoites using an ELISA technique (15). All female *Anopheles* mosquitoes captured in Walukuba and Kihihi were tested; in Nagongera testing was limited to 50 randomly selected female *Anopheles* mosquitoes per household per night due to the large numbers collected. Therefore, for each household and/or site it was possible to calculate multiple entomological metrics, including the average human biting rate (average number of *female Anopheles* mosquitoes caught in a household per day), the average sporozoite rate (the average proportion of mosquitos that tested positive for *Plasmodium falciparum*) and the entomological inoculation rate (EIR, the product of the household human biting rate and the site sporozoite rate).

### Statistical analyses

The purpose of these analyses was to model and quantify the development of immunity against symptomatic malaria, as a function of age and exposure, measured by the household EIR.

We modeled two specific types of immunity that have been previously described as components of immunity to malaria. We defined anti-parasite immunity as the ability to control parasite densities upon infection and anti-disease immunity as the ability to tolerate parasite infections without developing objective fever. Thus, for models of anti-parasite immunity, the outcome of interest was the parasite density recorded (using thick blood smear) at each parasite positive study visit. For models of anti-disease immunity the outcome of interest was the objective temperature recorded during parasite positive visits, conditional on the parasite density. In addition, we also modeled overall immunity against symptomatic malaria. For these analyses, the outcome of interest was the probability of presenting with fever given infection (parasite positivity).

In order to model the association between the outcomes and covariates of interest we used generalized additive models (gams). Gams provide a good framework, as they allow for smooth non-linear relationships. Details on the specific models explored are provided in the supplementary material. In summary, the models followed the following form.

1. Anti-parasite immunity

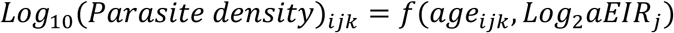
2. Anti-Disease immunity

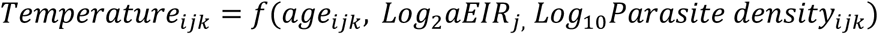
3. Overall immunity against symptomatic malaria

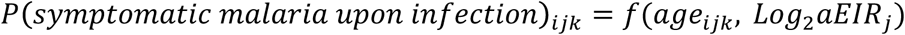

Where *i* is an index for individuals, *j* for households and *k* for specific visits. Thus, *age*_*ijk*_ represents the age of child *i* from household *j* during visit *k,* and *aEIR*_*j*_ represents the average annual EIR recorded for household *j.* We included the EIR as an average (time-invariant) covariate, as we were interested in modeling the impact of the average exposure to malaria over time on the development of clinical immunity.

All models were fitted in the R statistical framework using package mgcv(16). To account for clustering, all models included random effects at the individual and household levels. Best fitting models were selected based on Akaike’s Informaiton Criterion and the percent deviance explained.

## Acknowledgements

We thank all study participants who participated in this study and their families. We also thank hank the study team and the Makerere University–UCSF Research Collaboration and the Infectious Diseases Research Collaboration for administrative and technical support.

## Competing interests

None

## References

1. Bhatt S, Weiss DJ, Cameron E, Bisanzio D, Mappin B, Dalrymple U, et al. The effect of malaria control on Plasmodium falciparum in Africa between 2000 and 2015. Nature. Nature Publishing Group; 2015 Sep 16;526(7572):207–11.

2. World Health Organization (WHO). World malaria report 2016. Geneva: WHO. Embargoed until; 2016.

3. Filipe JAN, Riley EM, Drakeley CJ, Sutherland CJ, Ghani AC. Determination of the Processes Driving the Acquisition of Immunity to Malaria Using a Mathematical Transmission Model. PLoS Comput Biol. Public Library of Science; 2007 Dec 28;3(12):e255.

4. Smith TA, Leuenberger R, Lengeler C. Child mortality and malaria transmission intensity in Africa. Trends in Parasitology. 2001 Mar;17(3):145–9.

5. Snow RW, Omumbo JA, Lowe B, Molyneux CS, Obiero J-O, Palmer A, et al. Relation between severe malaria morbidity in children and level of Plasmodium falciparum transmission in Africa. The Lancet. 1997 Jun;349(9066):1650–4.

6. Griffin JT, Hollingsworth TD, Reyburn H, Drakeley CJ, Riley EM, Ghani AC. Gradual acquisition of immunity to severe malaria with increasing exposure. Proceedings of the Royal Society of London B: Biological Sciences. The Royal Society; 2015 Feb 22;282(1801):20142657.

7. Reyburn H, Mbatia R, Drakeley C, Bruce J, Carneiro I, Olomi R, et al. Association of transmission intensity and age with clinical manifestations and case fatality of severe Plasmodium falciparum malaria. JAMA. 2005 ed. 2005 Mar 23;293(12):1461–70.

8. Okiro EA, Al-Taiar A, Reyburn H, Idro R, Berkley JA, Snow RW. Age patterns of severe paediatric malaria and their relationship to Plasmodium falciparum transmission intensity. Malar J. 2009 ed. 2009;8:4.

9. Carneiro I, Roca-Feltrer A, Griffin JT, Smith L, Tanner M, Schellenberg JA, et al. Age-patterns of malaria vary with severity, transmission intensity and seasonality in sub-Saharan Africa: a systematic review and pooled analysis. PLoS ONE. 2010;5(2):e8988.

10. Idro R, Aloyo J, Mayende L, Bitarakwate E, John CC, Kivumbi GW. Severe malaria in children in areas with low, moderate and high transmission intensity in Uganda. Tropical medicine&international health : TM&IH. 2006 Jan;11(1):115–24.

11. Roca-Feltrer A, Carneiro I, Smith L, Schellenberg JR, Greenwood B, Schellenberg D. The age patterns of severe malaria syndromes in sub-Saharan Africa across a range of transmission intensities and seasonality settings. Malar J. 2010;9:282.

12. Rodríguez-Barraquer I, Arinaitwe E, Jagannathan P, Boyle MJ, Tappero J, Muhindo M, et al. Quantifying Heterogeneous Malaria Exposure and Clinical Protection in a Cohort of Ugandan Children. J Infect Dis. Oxford University Press; 2016 Oct 1;214(7):1072–80.

13. Struik SS, Riley EM. Does malaria suffer from lack of memory? Immunological Reviews. Munksgaard International Publishers; 2004 Oct 1;201(1):268–90.

14. Kamya MR, Arinaitwe E, Wanzira H, Katureebe A, Barusya C, Kigozi SP, et al. Malaria transmission, infection, and disease at three sites with varied transmission intensity in Uganda: implications for malaria control. Am J Trop Med Hyg. 2015 May;92(5):903–12.

15. Kilama M, Smith DL, Hutchinson R, Kigozi R, Yeka A, Lavoy G, et al. Estimating the annual entomological inoculation rate for Plasmodium falciparum transmitted by Anopheles gambiae s.l. using three sampling methods in three sites in Uganda. Malar J. 2014 Mar 21;13(1):111.

16. R Foundation for Statistical Computing. R:anguage and environment for statistical computing [Internet]. Vienna. Available from: http://www.R-project.org/

17. Lopera-Mesa TM, Doumbia S, Konaté D, Anderson JM, Doumbouya M, Keita AS, et al. Effect of red blood cell variants on childhood malaria in Mali: a prospective cohort study. The Lancet Haematology. Elsevier; 2015 Apr;2(4):e140–9.

18. Taylor SM, Parobek CM, Fairhurst RM. Haemoglobinopathies and the clinical epidemiology of malaria: a systematic review and meta-analysis. The Lancet Infectious Diseases. 2012 Jun;12(6):457–68.

19. Baird JK. Age dependent characteristics of protection v. susceptibility to Plasmodium falciparum. Annals of Tropical Medicine And Parasitology. Routledge, part of the Taylor&Francis Group; 1998 Jun 1;92(4):367–90.

20. Hviid L. Clinical disease, immunity and protection against Plasmodium falciparum malaria in populations living in endemic areas. Expert reviews in molecular medicine. Cambridge University Press; 1998 Jun 24;1998(04):1–10.

21. Bull PC, Lowe BS, Kortok M, Molyneux CS, Newbold CI, Marsh K. Parasite antigens on the infected red cell surface are targets for naturally acquired immunity to malaria. Nature medicine. Europe PMC Funders; 1998 Mar;4(3):358–60.

22. Wipasa J, Suphavilai C, Okell LC, Cook J, Corran PH, Thaikla K, et al. Long-Lived Antibody and B Cell Memory Responses to the Human Malaria Parasites, Plasmodium falciparum and Plasmodium vivax. Kazura JW, editor. PLoS Pathog. Public Library of Science; 2010 Feb 19;6(2):e1000770.

23. Langhorne J, Ndungu FM, Sponaas A-M, Marsh K. Immunity to malaria: more questions than answers. Nature Immunology. Nature Publishing Group; 2008 Jul 1;9(7):725–32.

24. Filipe JA, Riley EM, Drakeley CJ, Sutherland CJ, Ghani AC. Determination of the processes driving the acquisition of immunity to malaria using a mathematical transmission model. PLoS Comput Biol. 2007 Dec;3(12):e255.

25. Griffin JT, Hollingsworth TD, Reyburn H, Drakeley CJ, Riley EM, Ghani AC. Gradual acquisition of immunity to severe malaria with increasing exposure. Proceedings Biological sciences / The Royal Society. 2015 Feb 22;282(1801).

26. Griffin JT, Ferguson NM, Ghani AC. Estimates of the changing age-burden of Plasmodium falciparum malaria disease in sub-Saharan Africa. Nat Commun. Nature Publishing Group; 2014;5:3136.

